# Tirzepatide Synergizes with Leptin on Weight Loss and Restoring Metabolic Homeostasis in Diet-induced Obesity Model

**DOI:** 10.64898/2025.12.18.695152

**Authors:** Xun Sun, Yuexi Yin, Min Song, Brian A Droz, Yanzhu Lin, Kristen N. Beaty, Baohua Zhou, Hyun Cheol Roh, Jonathan M. Wilson, Paul F. Cain, Ricardo J. Samms, Patrick L. Sheets, Minrong Ai, Hongxia Ren

**Affiliations:** Herman B. Wells Center for Pediatric Research, Department of Pediatrics, Indiana University School of Medicine; Indianapolis, IN, USA; Center for Diabetes and Metabolic Diseases, Indiana University School of Medicine; Indianapolis, IN, USA; Stark Neurosciences Research Institute, Indiana University School of Medicine; Indianapolis, IN, USA; Lilly Research Laboratories, Eli Lilly and Company; Indianapolis, IN, USA; Department of Anatomy, Cell Biology & Physiology, Indiana University School of Medicine; Indianapolis, IN, USA

## Abstract

Leptin resistance limits anti-obesity efficacy. We identified a leptin-sensitizing mechanism through tirzepatide (TZP), a glucagon-like peptide-1 receptor (GLP-1R) and glucose-dependent insulinotropic polypeptide receptor (GIPR) dual-agonist. Our tirzepatide clinical trial revealed that circulating leptin levels at baseline correlated with weight loss efficacy in patients with obesity, suggesting leptin and tirzepatide could interact to achieve stronger effects on weight loss. Next, we utilized the diet-induced obesity (DIO) mice and demonstrated the synergistic effects of tirzepatide and leptin combination (TZP+Lep) on weight loss. TZP+Lep treatment further improved hepatic insulin sensitivity and upregulated thermogenetic gene expression in brown adipose tissue. Metabolic profiling under thermoneutrality revealed TZP+Lep treatment further reduced food intake and increased energy expenditure. Tirzepatide sensitized leptin signaling in hypothalamic pro-opiomelanocortin (POMC) and GLP-1R expressing neurons. TZP+Lep synergistically increased POMC neuronal firing by decreasing the inhibitory postsynaptic input. Together, our work showed combining tirzepatide and leptin as a potential way for better maintenance of metabolic homeostasis in obesity management.

**Highlights:**

- Tirzepatide and leptin synergistically promote weight loss through reduced food intake and increased energy expenditure.
- Tirzepatide and leptin synergistically improve insulin sensitivity and metabolic homeostasis with altered hepatic and adipose gene expression.
- Tirzepatide sensitizes leptin signaling in hypothalamic GLP-1R and POMC neurons.
- Tirzepatide and leptin synergistically increase POMC neuronal firing by decreasing inhibitory postsynaptic input.

## INTRODUCTION

Peptide hormones, including leptin produced by the adipose tissue, glucagon-like peptide-1 (GLP-1) and glucose-dependent insulinotropic polypeptide (GIP) produced by gastrointestinal tract, relay peripheral adiposity and nutritional signals to central nervous system to regulate feeding and glucose metabolism (*1, 2*). Deciphering the neurohormonal functions of these hormones will uncover therapeutic interventions to combat obesity epidemic by achieving more weight loss efficacy and mitigating recidivism. GLP-1 is an incretin hormone secreted from the enteroendocrine L cells in response to meal ingestion to promote insulin secretion and reduce feeding (*3*). GLP-1 exerts its biological actions through the cognate G protein-coupled receptor, GLP-1R, in the central and peripheral tissues (*4*). GLP-1R agonist injectables, including liraglutide, and semaglutide, led to significant and meaningful weight loss and were FDA approved obesity therapies (*5*). GIP is another incretin hormone secreted from K cells of the proximal GI tract in response to food ingestion. Tirzepatide, a GLP-1 and GIP receptor dual agonist, was recently approved for obesity treatment (*6*). Tirzepatide treatment resulted in substantial weight reductions in patients with obesity (*7*), patients with obesity and diabetes (*8*), and patients living with obesity after lifestyle intervention (*9*). Tirzepatide achieved greater weight loss than semaglutide in head-to-head trials involving people with obesity with and without diabetes (*10, 11*). Despite the increasing use of these highly effective obesity management medications, patients still experienced significant weight regains after treatment cessation(*12*).

Leptin is an adipose-derived hormone and often tightly correlates with fat mass (*13*). It reflects long-term energy storage in organisms and regulates energy homeostasis by modulating feeding behavior and thermogenesis (*14*). Leptin binds to the long-form leptin receptor (OB-Rb, LepR hereafter), which is highly expressed in the hypothalamus, and signals through JAK-STAT pathway (*1, 15*). Hyperleptinemia, characterized by elevated leptin levels, is commonly observed in individuals who are obese or overweight. As a result, leptin resistance presents a significant challenge in developing leptin as an effective monotherapy for obesity.

Preclinical studies using the combination of leptin and other hormones have provided insights into potential new approaches for obesity treatment (*16*). For instance, a recent study demonstrated the efficacy of the engineered GLP-1R/LepR dual agonist in reducing food intake and body weight in rodent models (*17*). However, the synergistic effect of the GLP-1 and leptin on energy balance and peripheral metabolism is unclear.

We observed that higher baseline circulating leptin levels correlate with greater weight loss in a clinical trial of female and male persons with obesity under various doses of tirzepatide treatment. We hypothesized that leptin and GLP-1 will synergize on weight loss and improve energy metabolism. In this study, we determined the effects of tirzepatide and leptin combination treatment on weight loss and feeding in diet-induced obesity (DIO) mouse model. We characterized peripheral metabolism and compared the metabolic profiles of the combination treatment versus single peptide hormone treatment under thermoneutrality using indirect calorimetry. We performed mechanistic studies to investigate the effects of combination treatment on leptin signaling and neurological functions using immunohistochemistry and electrophysiological approaches, which provides mechanistic depth lacking in prior translational studies. Together, our work presents integrative physiology advance that links clinical data, systemic metabolism, and neural circuitry.

## RESEARCH DESIGN AND METHODS

### Clinical trial design and data analysis

This phase 2b clinical trial design was described previously (*19*). Briefly, in this a double-blind (patient and investigator) study, participants (aged 18-75 years) had type 2 diabetes for 6 months or longer (HbA1c 7.0%-10.5%), which was inadequately controlled with diet and exercise alone or with stable metformin therapy, and a body mass index of 23-50 kg/m^2^ (*18*). Overall, 318 participants were randomized (1:1:1:1:1:1) to receive once-weekly subcutaneous 1 (N = 53), 5 (N = 55), 10 (N = 52), or 15 mg (N = 53) tirzepatide, 1.5 mg dulaglutide (N = 54), or placebo (N = 51), with 316 participants included in the modified intention-to-treat population. Participants were treated for 26 weeks after a 1-week screening and 2-week lead-in period. The study (NCT03131687) was approved by the relevant ethics committees and conducted in accordance with the principles of the Declaration of Helsinki, Council of International Organizations of Medical Sciences International Ethical Guidelines, and Good Clinical Practice guidelines. All participants provided written informed consent prior to participation. Biomarkers were quantified in serum or EDTA plasma collected in the fasting state from patients at baseline, 4, 12, and 26 weeks and stored at −80°C until analysis. Leptin (R&D Systems, Minneapolis, MN) was measured by immunoassay.

Analyses were performed on the modified intention-to-treat population, excluding data from the dulaglutide group and data after study drug discontinuation or rescue drug initiation. Subjects included in analyses had non-missing baseline values and at least one non-missing postbaseline value of the response variable. Subjects were divided into subgroups based on the median value of leptin within each sex. Mixed model with repeated measures (MMRMs) were used to analyze changes in body weight within each subgroup, with baseline body weight included as a covariate.

### Mouse models

Diet-induced obese (DIO) C57/BL6 male mice (#DIO-B6-M) were purchased from the Taconic. POMC-EGFP transgenic mice (#009593) were purchased from the Jackson Laboratory (JAX). The animals were randomly housed under a 12-hour light/dark cycle, with five mice per cage or individually, depending on the experimental requirements. Mice were fed either a normal chow diet (NCD) (#5053, LabDiet, St. Louis, MO) or a high-fat diet (HFD) (#D12492 rodent diet 60kcal% fat, Research Diets, New Brunswick, NJ) with access to water. All animal procedures are followed and approved by Indiana University School of Medicine Animal Care and Use Committee.

### Peptides and recombinant proteins

Tirzepatide (LY3298176) was previously described (*6*). Recombinant leptin protein (LY355101) used in Fig. 5B-E, Fig. 6E-G and Fig. 7 was previously described (*48*). Fc-Leptin (FcLep) used in this study is a recombinant protein with a mouse immunoglobulin gamma (IgG) constant domain (Fc) fused to the N-terminus of human leptin. Briefly, DNA construct encoding mouse IgG1-Fc, a linker amino acid sequence of GGGGS and a human leptin protein lacking its signal peptide was generated. Recombinant FcLep was purified following the standard Protein-G based chromatography.

### In vivo efficacy studies

In vivo efficacy experiments were conducted in 40-week DIO C57/BL6 male mice. Before the acclimation, the diet was switched from HFD to NCD. Mice were randomly grouped into 4 groups and single-housed (vehicle = 6, FcLep = 6, TZP = 9, TZP+FcLep = 9). After 5 days of acclimation and 3 days of daily subcutaneous injections with 40 nM Tris buffer (pH = 8), daily dose injection started from day 0 and lasted three weeks. The doses of tirzepatide and FcLep were 3 nmol/kg and 40 nmol/kg, respectively. Compounds were co-formulated and made fresh weekly. The daily body weight measurement, food intake measurement, and injection were done at 3 pm-4 pm. Of note, two mice in the combination group shredded food and their food intake data were removed from the data set accordingly. The food intake data on day 4 was missed due to the erroneous operation of the animal technician. The average food intake reduction was calculated from the average food intake under-dosing treatment minus the average food intake under-saline treatment. Body composition was determined by Nuclear Magnetic Resonance (NMR) Relaxometry, using EchoMRI on days −1 and 13. The percentage of food intake reduction and body weight reduction during the third week of dosing was calculated based on the data on Day 14. After 21 days of dosing administration, mice were fasted overnight and refed at 9 AM on day 22. Feeding activity was monitored for the next 24 hours. Mice were treated with a dose injection at 9 am on day 23 and euthanized 2 hours later for tissue collection.

### Metabolic analyses

Metabolic analyses were conducted in 22-week DIO C57/BL6 male mice with an average body weight of 53 g. Mice were randomly grouped into 4 groups (vehicle, FcLep, tirzepatide, TZP+FcLep, n = 6 per group) and single-housed under thermoneutral condition (∼28°C). Prior to measurements, mice were switched to a normal chow diet (NCD) and acclimatized to the metabolic cages for 7 days. All mice were pre-dosed with vehicle for 3 days followed by 3 weeks dosing treatment. Metabolic parameters were monitored and recorded continuously using the TSE calorimetric system (TSE System, Germany). Designating day 0 is the first dosing day, mouse body composition was conducted on day −3 and day 21 using nuclear magnetic resonance (NMR) system (Bruckner Minispec mq10; Bruker Corporation, Billerica, MA, USA).

### Insulin levels measurement

Blood samples were collected by cardiac puncture. Serum insulin concentrations were determined with an enzyme-linked immunosorbent assay (ELISA) kit (#10-1247-10; Mercodia, Uppsala, Sweden).

### Quantitative reverse transcription polymerase chain reaction (qRT-PCR)

The total RNA of the hypothalamus and hindbrain was extracted with TRIzol Reagent, while the RNA of brown adipose tissue and liver was extracted by RNeasy Plus Mini Kit (#74136; Qiagen, Hilden, Germany). Superscript II reverse transcriptase (#18064014; Invitrogen) was used to synthesize template cDNA, and the specific productions of quantitative real-time PCR (qRT-PCR) were validated by melting curve, which was shown by CFX Connect Real-Time system (Bio-Rad, Hercules, CA). Primer sequences can be found in Table S1.

### Combined immunohistochemistry and RNAscope following compounds dosing

Male DIO mice were divided into 6 treatment groups (n = 8/group). The mice were changed to regular chow 1 week before the first dosing. Food intake and body weight of the mice were registered from study day −3 to day 6. Group 5 and 6 were pair-fed to group 4. The mice received daily subcutaneous doses with tirzepatide (3 nmol/kg) or vehicle (40 nM Tris buffer) for 6 days. 16 hours before the last dose, food was removed, and 1 hour after the last dose, the mice received an intraperitoneal dose of recombinant leptin (LY355101) or vehicle. After 45 minutes, the mice were perfused with neutral-buffered formalin, and the brains were dissected. Formalin-fixed paraffin-embedded brain sections (3 sections per mouse per triplex) underwent RNAscope in situ RNA hybridization (ISH) for the detection of Lepr (ACDBio probe #418858-C2), Glp-1r (ACDBio probe #418858-C2), Gipr (ACDBio probe #319128-C1), Pomc (ACDBio probe #314088-C2) and Agrp (ACDBio probe #400718-C1), followed by immunofluorescence (IF) staining for pSTAT3 protein (Cell Signaling anti-pSTAT3 #9145).

After ISH and IF, sections were counterstained with DAPI, cover-slipped with a fluorescence mounting medium. Slides were scanned under 20x objective in an Olumpus VS-120 slide scanner and imported into VIS software (Visopharm, Denmark). Cells expressing mRNA of interest were counted with reference to DAPI. Cells co-expressing multiple mRNA of interest and/or pSTAT3 protein were defined and quantified by custom written add-on applications for Visopharm software.

### In vitro pSTAT3 response to leptin treatment

HEK293 cells were transiently transfected with control DNA construct (pcDNA3.1) or DNA constructs expressing human GLP1R, human LEPR or both following standard Fugene® method. Transfected cells were seeded in 96-well plate at ∼60,000 cells per well and cultured overnight. Cells were then treated with increasing concentrations of leptin (LY355101) in the absence or presence of EX4 (1nM and 10nM, respectively) at 37 °C for 15 minutes. Phosphorylated STAT3 (pSTAT3) was then measured with an AlphaLISA Sure Fire Ultra kit (Revvity, #ALSU-PST3-A10K) following manufacture’s instruction. The pSTAT3 dose-response curves were generated with four parameter non-linear curve fit using GraphPad Prism software.

### Acute brain slice preparation

Experiments were performed on 6 to 8-week-old POMC-EGFP transgenic mice. Brains were dissected and transferred in chilled carbogenated choline (in mM: 25 NaHCO3, 1.25 NaH2PO4, 2.5 KCl, 0.5 CaCl2, 7 MgCl2, 25 D-glucose, 110 C5H14CINO, 11.6 C6H7NaO6, 3.09 C3H3NaO3). Coronal brain slices were sectioned in 300 µm using a vibratome (VT1200s Leica, Wetzlar, Germany). Slices were transferred to carbogenated artificial cerebrospinal fluid (aCSF) solution (in mM: 127 NaCl, 25 NaHCO3, 1.25 NaH2PO4, 2.5 KCl, 25 D-glucose, 2 CaCl2, 1 MgCl2) at 37°C for 30 minutes and then at room temperature for at least 45 minutes before recording.

### Electrophysiology

We measured the electrophysiology of the POMC neurons in the mouse ARH. The POMC neurons will be identified by the green fluorescence protein (GFP), a method we previously used (*49*). The POMC-EGFP transgenic mice have EGFP expression directed to POMC-expressing neurons by the mouse Pomc promoter/enhancer regions (https://www.jax.org/strain/009593). Recording pipettes were fabricated from borosilicate capillaries using a horizontal puller (P-97; Sutter Instrument, Novato, CA) with series resistance between 2-4 MΩ. Fluorescently labeled cells were visualized using an LED illumination system (CoolLED pE-4000, UK). For intrinsic spontaneous action potential firing recordings, a potassium-based internal solution was used (in mM: 128 K-gluconate, 10 HEPES, 10 sodium phosphocreatine, 4 MgCl2, 4 sodium ATP, 0.4 sodium GTP, 3 ascorbic acid, 1 EGTA, 4 mg/mL biocytin). For intrinsic excitatory injecting current steps recordings, same potassium-based internal solution was used. NMDA receptor antagonist CPP (5 µM), AMPA receptor antagonist NBQX (5 µM), and GABAA receptor antagonist GABAzine (10 µM) were added to the aCSF bath solution. For spontaneous inhibitory postsynaptic current (sIPSC) recordings, pipettes were filled with high chloride cesium-based internal solutions (in mM: 120 CsCl, 4 MgCl2, 10 HEPES, 10 EGTA, 2 magnesium ATP, 0.5 sodium GTP, 5 Lidocaine). CPP and NBQX were added to the aCSF bath solution. Whole-cell patch clamp recordings were performed targeting cells that were 50-80 µm deep in the slice and the aCSF solution was maintained at 30-31 °C. Cells with series resistance higher than 35 MΩ were excluded. Current-clamp recordings were bridge balanced. Recordings were filtered at 4 kHz for intrinsic recordings and at 2 kHz for sIPSC recordings and were digitized at 10 kHz using Multiclamp 700B amplifier (Molecular Devices, San Jose, CA). Cells were separated into two groups: one treated with Lep (20 µM) first, and the other with TZP (10 µM) first. A baseline was recorded for 6 minutes before the application of the first drug. Following 9 minutes of recording post the first drug application, the second drug was added to the bath. Cells were then recorded for an additional 9 minutes. Data was calculated using the last 3 minutes of each recording session. Custom MATLAB (The MathWorks, Natick, MA) routines were used to process electrophysiology data offline.

### Quantification and statistical analysis

Statistical significance was determined by Student’s t-test, one-way, or two-way ANOVA where appropriate using GraphPad Prime software (San Diego, CA). Group data were presented as mean and error bars indicated the SEM. The statistical significance was defined as *p ≤ 0.05, **p ≤ 0.01, ***p ≤ 0.001, and ****p ≤ 0.0001.

### Data and Resource Availability

The data sets generated during the study are available from the corresponding author upon reasonable request.

## RESULTS

### Circulating leptin level correlates with weight loss percentage in patients undergoing tirzepatide treatment

To understand the relationship of circulating leptin level and the effect of tirzepatide treatment, we conducted a post hoc analysis of a phase 2 trial where patients living with diabetes and obesity received placebo or tirzepatide (1mg, 5mg, 10mg, 15mg) treatments (*18*). Leptin level and body weight over the treatment period (0, 4, 8, 12, 16, 20 and 26 weeks) (*19*) were used for sub-group data analysis. At baseline (week 0), the median plasma leptin levels were 29.3 ng/ml and 10.5 ng/ml for female and male subjects, respectively. Subsequently, we analyzed the body weight change over trial period from four participant groups: females with baseline leptin above 29.3ng/ml (female^High-Lep^, N=53) and below 29.3ng/ml (female^Low-Lep^, N=53), males with baseline leptin above 10.5ng/ml (male^High-Lep^, N=59) and below 10.5ng/ml (male^Low-Lep^, N=58). At the end of 26-week trial, the percent body weight change from baseline in female^High-Lep^ patients receiving 10mg and 15mg TZP was −12.52 ± 2.06% (average ± standard error) and −13.78 ± 1.76%, respectively (Fig. 1A). In contrast, female^Low-Lep^ patients receiving 10mg and 15mg TZP had −10.49 ± 1.67% and −9.16 ± 1.36% body weight change, respectively (Fig. 1B). In male^High-Lep^ patients receiving 10mg and 15mg TZP, the percentage body weight change from baseline was −8.51 ± 1.76% and −11.77 ± 2.12%, respectively (Fig. 1C). In contrast, male^Low-Lep^ patients receiving 10mg and 15mg TZP had −6.61 ± 1.22% and −6.84 ± 1.93% body weight change, respectively (Fig. 1D). Therefore, higher baseline leptin level seems to predict the outcome of weight loss efficacy of subsequent TZP treatment for both male and female patients. Based on these data, we hypothesize that co-administering tirzepatide and leptin may result in greater weight loss compared to tirzepatide alone.

**Fig. 1.**
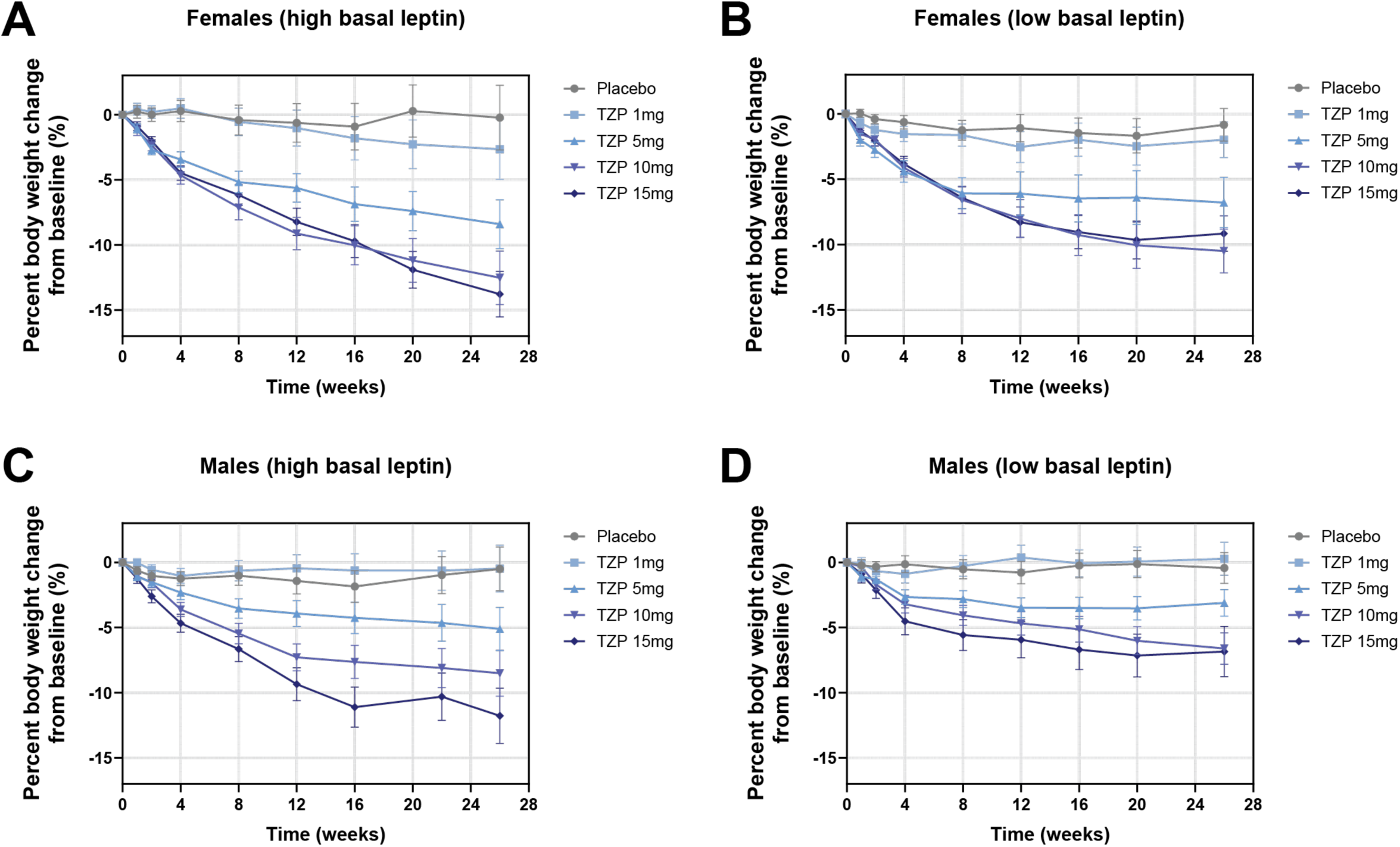
The correlation of basal leptin levels and body weight change under tirzepatide treatment. **(A-B)** Percentage body weight change under placebo or TZP treatment (1, 5, 10, 15 mg) in female patients with high (A) or low (B) basal leptin. **(C-D)** Percentage body weight change under placebo or TZP treatment (1, 5, 10, 15 mg) in male patients with high (C) or low (D) basal leptin. In all participants receiving placebo and various doses of tirzepatide treatments, the median baseline leptin is 29.3 ng/ml and 10.5 ng/ml for female and male participants, respectively. N=53 for female^High-Lep^, N=53 for female^Low-Lep^, N=59 for male^High-Lep^, N=58 for male^Low-Lep^.

### The synergistic effect of tirzepatide and leptin on weight loss in mice

To evaluate the combination effect of tirzepatide and leptin, the DIO mice received daily subcutaneous injections of tirzepatide (TZP, 3 nmol/kg), a mouse IgG1-Fc and human leptin fusion protein (FcLep see methods, 40 nmol/kg), or a combination of tirzepatide and leptin (TZP+FcLep) for three weeks, following 5-day diet switch acclimation and 3 daily vehicle (40 nM Tris, pH 8) injection acclimation (Fig. 2A). The body weight of the FcLep group remained comparable to vehicle control due to leptin resistance. In contrast, both TZP and TZP+FcLep groups exhibited significant reductions in body weight from day 1 to day 21 compared with vehicle or FcLep groups. Notably, a significant difference between TZP and TZP+FcLep groups emerged from day 12, indicating a synergistic effect of tirzepatide and leptin (Fig. 2B and Fig. S1A). Additionally, the TZP+FcLep group showed a moderate yet significant reduction in body weight than TZP alone within the first two days of dosing, suggesting that the synergistic effect was independent of weight loss (Fig. 2C).

**Fig. 2.**
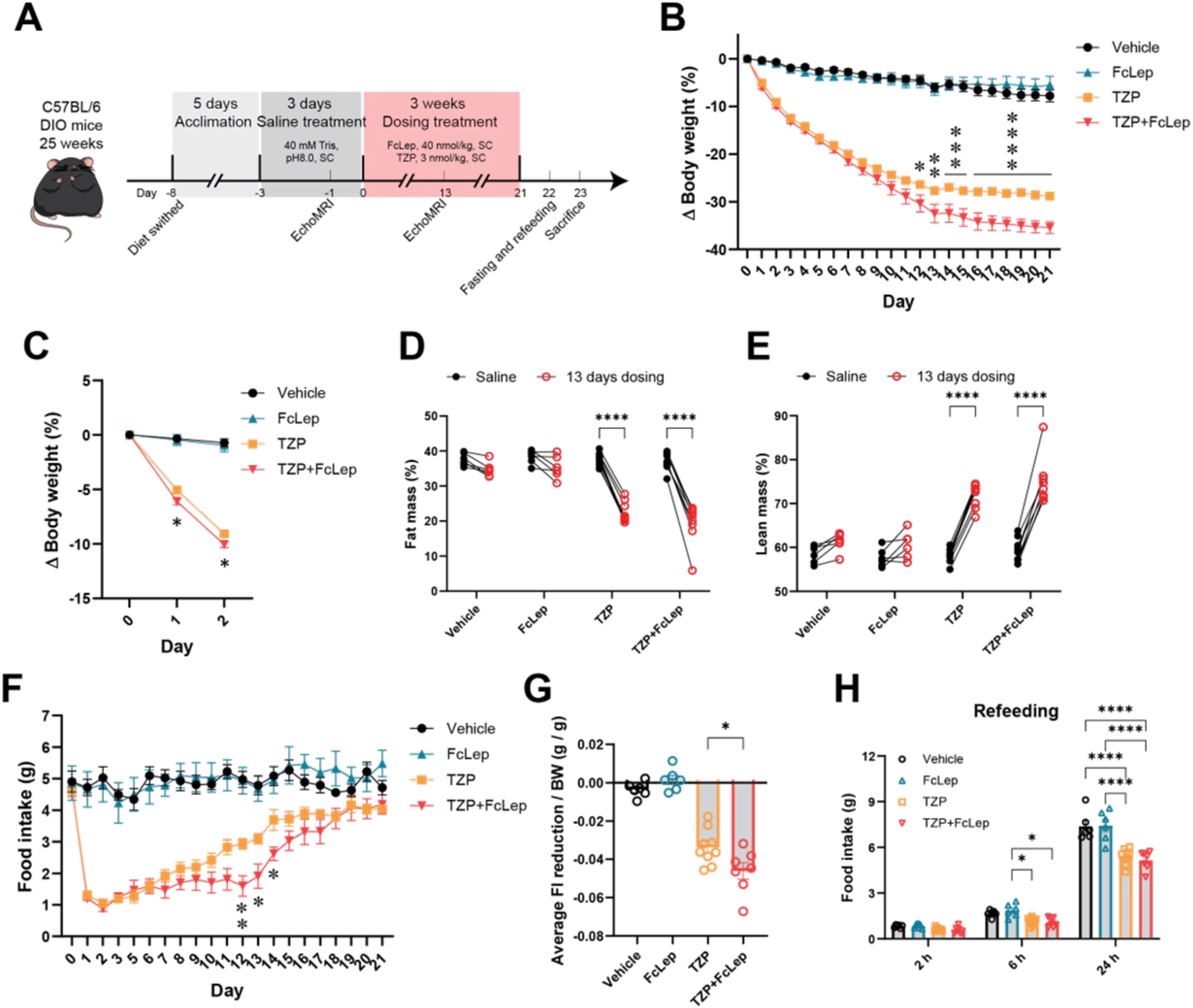
Tirzepatide synergized with leptin on weight loss. Fc-leptin = mouse IgG1-Fc and human leptin fusion recombinant protein, TZP = tirzepatide, SC = subcutaneous, FI = food intake, BW = body weight. (A) Schematic illustration of the study design (4 dosing groups: vehicle, FcLep, TZP, TZP+FcLep). (B) Normalized body weight reduction (%) throughout 21 days of dosing. Two-way ANOVA. * presented the comparison between TZP and TZP+FcLep. (C) Normalized body weight reduction (%) in the first 3 days. Two-way ANOVA. * presented the comparison between TZP and TZP+FcLep. (D-E) Fat mass and lean mass (% of body weight) on days −1 and day 13. Two-way ANOVA. (F) Daily food intake measured throughout 21 days of dosing. Two-way ANOVA. * presented the comparison between TZP and TZP+FcLep. (G) Average 24 h food intake reduction (normalized to body weight). One-way ANOVA.(H) Food intake in 2, 6, and 24 h after overnight fasting. Two-way ANOVA. n = 6-9 per group. *p < 0.05, ^∗∗^p < 0.01, ^∗∗∗^p < 0.001, ^∗∗∗∗^p < 0.0001. Data are displayed as means ± SEM.

Tirzepatide decreases weight primarily through reduced appetite, energy intake, and fat mass (*20*). We next investigated the metabolic changes undergoing tirzepatide and leptin co-administration. The body composition was assessed one day before and after 13 days of dosing treatment. Both TZP and TZP+FcLep groups showed a significant reduction in fat mass percentage, with decreases of 15-17% (Fig. 2D and Fig. S1B). Concurrently, an increase in the lean mass percentage was observed, with gains of 13% in the TZP group and 15% in the TZP+FcLep group (Fig. 2E and Fig. S1C). The DIO mice showed a notable reduction in food intake after receiving TZP or TZP+FcLep treatment. A significant difference was observed between these two groups from day 12 to day 14, followed by a compensatory phase (Fig. 2F). Although no significant difference in cumulative food intake was detected between TZP and TZP+FcLep groups (Fig. S1D), the TZP+FcLep group exhibited a further reduction of average daily food intake compared with TZP alone (Fig. 2G and Fig. S1E). Moreover, both TZP and TZP+FcLep groups reduced rebound food intake following an overnight fast on day 22, suggesting a reduction in rebound hyperphagia (Fig. 2H). These data indicate that tirzepatide, in synergy with leptin, enhances the weight loss effect by reducing food intake and fat mass. As noted, the difference in food intake between the TZP and TZP+FcLep groups gradually diminished during the final week of dosing, suggesting a compensatory phase. Indeed, all the mice in the TZP+FcLep group showed an increase in food intake (Fig. S1F). Interestingly, only one of the nine mice gained body weight, while the others experienced further weight loss (Fig. S1G & H). This suggests that additional factors may contribute to potentiating weight loss during the compensatory phase.

### Improved insulin sensitivity and metabolic gene expression profile in DIO mice

Insulin resistance is a crucial feature of metabolic syndrome triggered by DIO. Both TZP and TZP+FcLep groups exhibited a marked reduction in serum insulin levels under ad libitum feeding conditions, indicating improved insulin resistance (Fig. 3A). In line with the weight loss, the weights of epididymal white adipose tissue (EWAT), brown adipose tissue (BAT), and liver in TZP and TZP+FcLep groups were all significantly lower compared to vehicle or FcLep groups (Fig. 3B-D and Fig. S2A-C). Although these tissue weights did not differ significantly between these two groups, the TZP+FcLep group always had a lower value than the TZP group. Moreover, a noticeably darker BAT color in the TZP+FcLep group was observed upon dissection. The qPCR results show that the TZP+FcLep group had the highest Irs2 level, a hepatic gene critical for preventing insulin resistance, supporting the idea of enhanced hepatic insulin sensitivity (Fig. 3E).

**Fig. 3.**
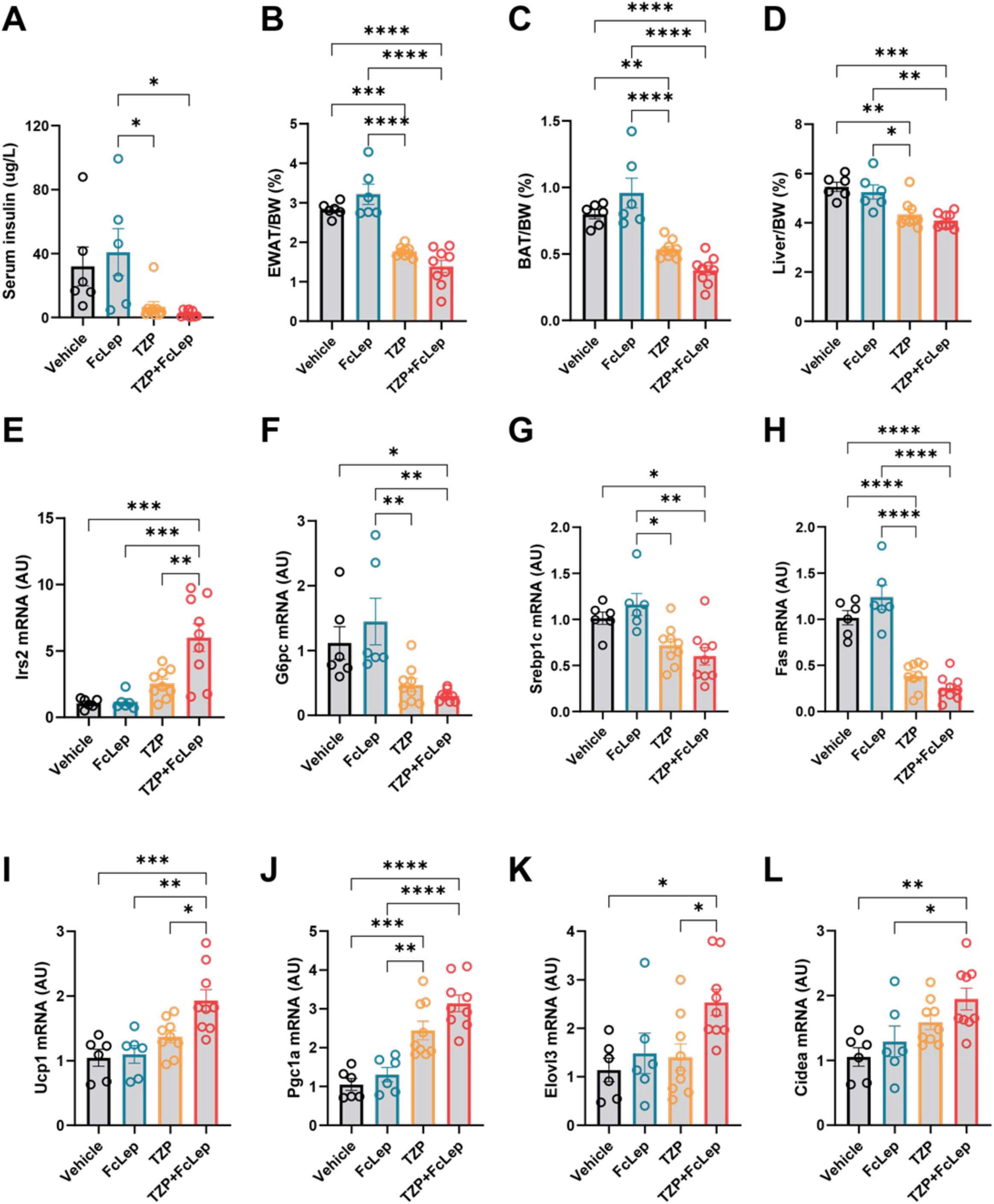
Improved hepatic insulin sensitivity and altered metabolic genes under TZP+FcLep treatment. EWAT = epididymal white adipose tissue, BAT = brown adipose tissue, BW = body weight. (A) Serum insulin level under ad libitum condition. (B-D) Percentage of EWAT, BAT, and liver weights normalized by body weight. (E-H) qPCR analysis of Irs2, G6pc, Srebp1c, and Fas in liver. (I-L) qPCR analysis of Ucp1, Pgc1a, Elovl3, and Cidea in BAT. One way ANOVA was performed. n = 6-9 per group. *p < 0.05, ^∗∗^p < 0.01, ^∗∗∗^p < 0.001, ^∗∗∗∗^p < 0.0001. Data are displayed as means ± SEM.

We further examined the mRNA expression of hepatic genes involved in glucose homeostasis and lipid metabolism. Both TZP and TZP+FcLep groups showed downregulation of G6pc, Gck, Pklr, and Pdk4, which may indicate improved glucose tolerance (Fig. 3F and Fig. S2D-F). No change was observed in Chrebp, a master regulator associated with glycolysis and de novo lipogenesis (Fig. S2G). Additionally, genes involved in fatty acid synthesis and lipid metabolism, including Srebp1c, Fas, Acc1, and Scd1, were downregulated in both TZP and TZP+FcLep groups (Fig. 3G-H and Fig. S2H-I). Moreover, we examined the expression of thermogenic and mitochondrial genes in BAT. The mRNA expression analysis revealed that the TZP+FcLep group, compared to the other three groups, expressed higher levels of Ucp1, Pgc1a, Elovl3, and Cidea, while expressing comparable levels of Prdm16, Cox7a1, and Cox8b, indicating increased energy expenditure (Fig. 3I-L and Fig. S2J-L). These results suggest that tirzepatide and leptin work synergistically to improve metabolic homeostasis, as evidenced by enhanced insulin sensitivity, decreased gluconeogenesis, and increased lipolysis and thermogenesis.

### Tirzepatide synergized with leptin on weight loss by increasing energy expenditure

We next examined metabolic parameters following 3 weeks of treatment using indirect calorimetry conducted under thermoneutrality condition (Fig. 4A). TZP+FcLep group again showed significant weight loss and feeding suppression compared to TZP alone (Fig. 4B and C). Both TZP and TZP+FcLep groups increased lean mass percentage and decreased fat mass percentage (Fig. 4D and E). Notably, the TZP+FcLep group demonstrated a greater overall change in body composition (Fig. 4F). Raw values of body weight, food intake, and body composition are provided in Supplemental Fig. 3A-E. Indirect calorimetry revealed that TZP+FcLep significantly increased oxygen consumption (VO_2_) and energy expenditure compared to TZP alone during the second week, particularly on day 10 and day 11. By the third week, both TZP and TZP+FcLep groups exhibited lower VO_2_ and energy expenditure than vehicle and FcLep groups, likely due to reduced body weight (Fig. 4G and Fig. S3F). Individual energy expenditure data were plotted against body weight. Beginning in the second week, a clear separation emerged between the TZP-treatment groups and the vehicle and FcLep groups, indicating a TZP-driven effect. Notably, the TZP+FcLep group exhibited higher energy expenditure compared to the TZP group at equivalent body weights after the first week of treatment (Fig. 4H). A decrease of respiratory exchange ratio (RER) was observed in both TZP and TZP+FcLep groups during the first two weeks and gradually increased in the third week, suggesting a preferential shift toward the lipid utilization as the primary energy source in the initial treatment period and a compensation phase consistent with feeding rebound (Fig. 4I). No significant differences in locomotor activity were detected among the four groups (Fig. S3G).

**Fig. 4.**
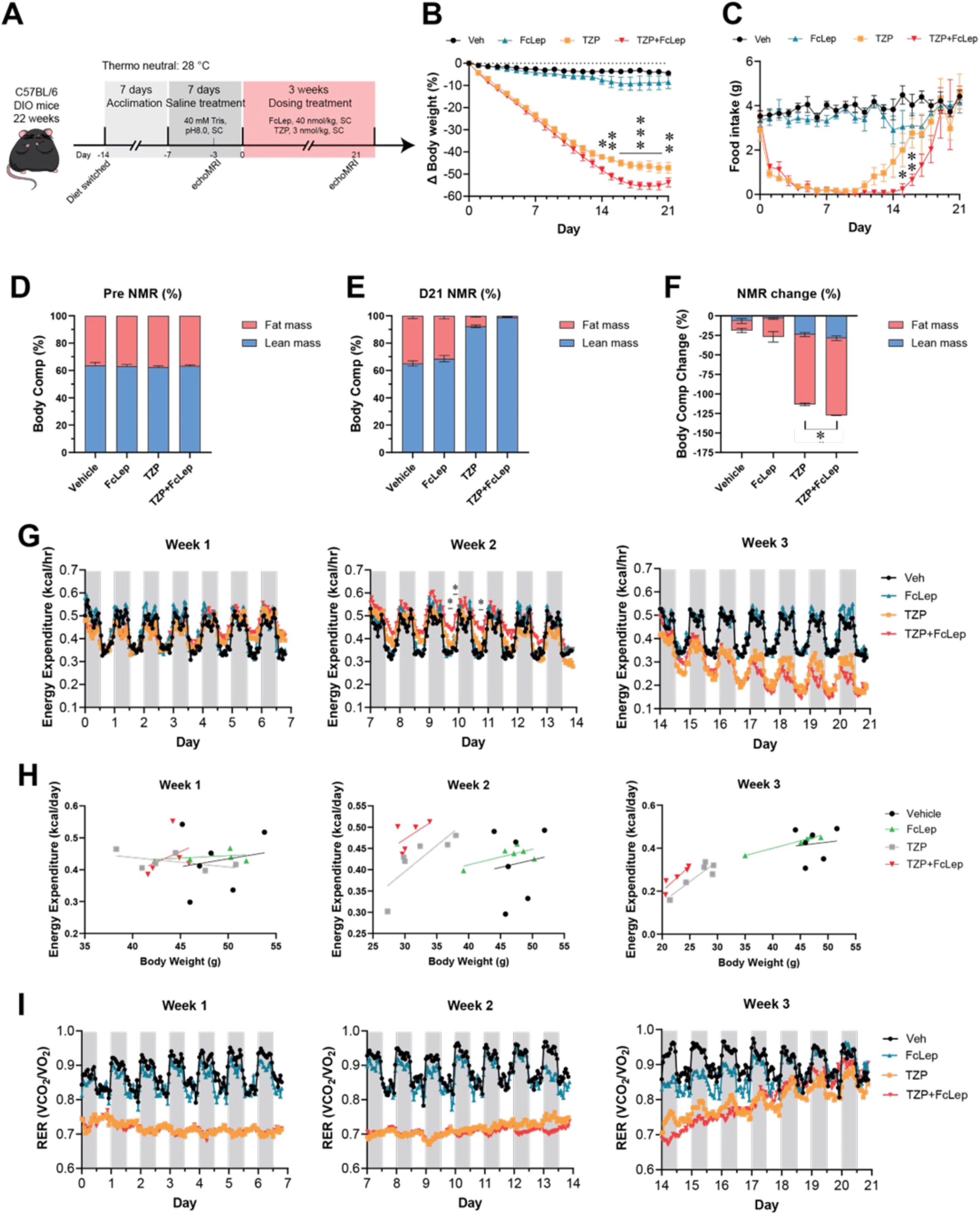
Increased energy expenditure and oxygen consumption in DIO mice treated with TZP+FcLep compared to TZP alone. SC = subcutaneous, NMR = nuclear magnetic resonance, VO_2_ = oxygen consumption, RER = respiratory exchange ratio. (A) Schematic illustration of the study design (4 dosing groups: vehicle, FcLep, TZP, TZP+FcLep). n = 6 per group. (B) Normalized body weight reduction (%) throughout 21 days of dosing at thermoneutrality. Two-way ANOVA. * presented the comparison between TZP and TZP+FcLep. (C) Daily food intake measured throughout 21 days of dosing at thermoneutrality. Two-way ANOVA. * presented the comparison between TZP and TZP+FcLep. (D-E) Fat mass and lean mass (%) of DIO mice before 3-day and after 21-day of dosing treatment. (F) Percentage change of fat mass and lean mass after treatment. One-way ANOVA. (G) Rhythms energy expenditure of DIO mice during 3-week treatment. Unpaired two-tailed Student’s t test was performed. *p < 0.05, statistically significant at three consecutive time points, compared to TZP. (H) Scatter plot of average daily energy expenditure and average daily body weight during each week of treatment. (I) Respiratory exchange ratio during 3-week treatment. n = 5-6 per group. *p < 0.05, ^∗∗^p < 0.01, ^∗∗∗^p < 0.001, ^∗∗∗∗^p < 0.0001. Data are displayed as means ± SEM.

### Tirzepatide sensitized hypothalamic leptin signaling

We aimed to investigate how tirzepatide synergizes with leptin to enhance energy homeostasis and suppress feeding. To begin, we assessed the sensitivity of LepR signal pathway in the hypothalamus by immuno-detecting phosphorylated STAT3 (pSTAT3) in mice treated with vehicle, TZP, or pair-feeding (PF) (Fig. 5A). In situ hybridization together with immunohistochemistry showed the expression of pSTAT3 in POMC, Agouti-related peptide (AgRP), and GLP-1R neurons in ARH (Fig. 5B and C). Both the TZP and PF groups exhibited a significant increase in the proportion of pSTAT3+ cells among POMC neurons, suggesting an enhanced LepR signaling in POMC neurons (Fig. 5D). A similar trend was observed in AgRP neurons (Fig. S4). Importantly, TZP—but not PF—significantly increased the responsiveness of GLP-1R-expressing neurons to exogenous leptin, indicating a direct sensitization of leptin signaling in these cells (Fig. 5E). The GIPR mRNA signal in the mediobasal hypothalamus is minimal (Fig. 5C), preventing accurate quantification.

**Fig. 5.**
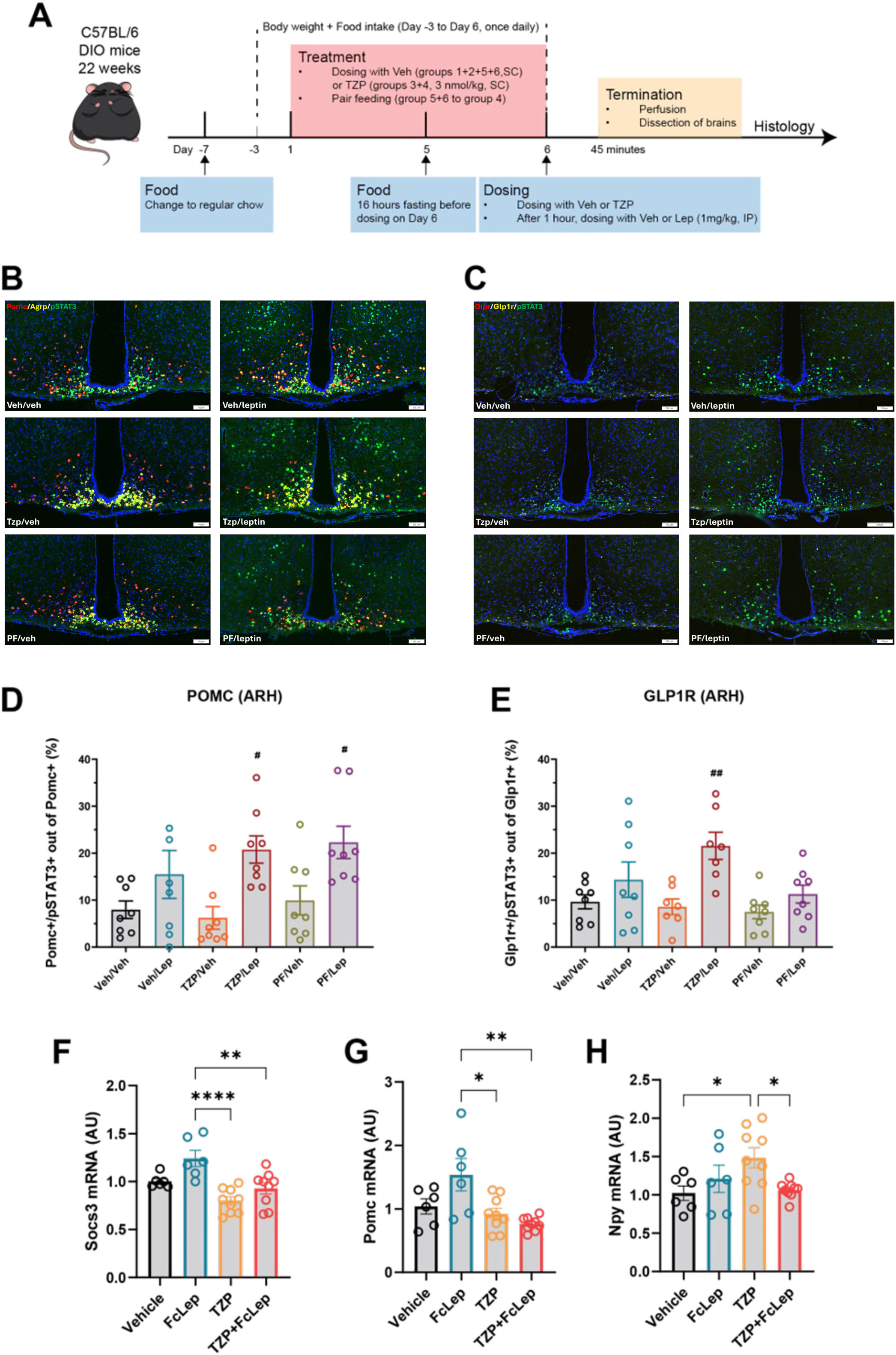
Tirzepatide sensitizes leptin signaling in the brain. Veh = vehicle, Lep = leptin (LY355101), TZP = tirzepatide, PF = pair-feeding, SC = subcutaneous, POMC = pro-opiomelanocortin, ARH = arcuate nucleus of hypothalamus, GLP1-R = glucagon-like peptide-1 receptor, pSTAT3 = phosphorylated STAT3. (A) Schematic illustration of the study design (6 dosing groups: Veh + Veh, Veh + Lep, TZP + Veh, TZP + Lep, PF + Veh, PF + Lep). n = 8 per group. (B) Representative images of Agrp and Pomc positive cells co-expressing pSTAT3 protein. (C) Representative images of Glp1r+, Gipr+ cells and pSTAT3 protein expression. (D) Percentage of Pomc+/pSTAT3+ cells out of all Pomc+ cells in ARH, calculated as the average of 3 sections. n = 7-8. Dunnett’s test one-factor linear model. ^#^P < 0.05 compared to TZP/Veh. (E) Percentage of Glp1r+/pSTAT3+ cells out of all Glp1r+ cells in ARH, calculated as the average of 3 sections. n = 7-8. Dunnett’s test one-factor linear model. ^##^P < 0.01 compared to TZP/Veh. (F-H) qPCR analysis of Socs3, Pomc, and Npy in the hypothalamus from DIO mice after 3 weeks of reagent treatment. n = 6-9 per group. One-way ANOVA. *p < 0.05, ^∗∗^p < 0.01, ^∗∗∗∗^p < 0.0001. Data are displayed as means ± SEM.

To determine that the increased energy expenditure and satiety shown in Fig. 2 and Fig. 3 were linked to enhanced leptin sensitivity, we examined the gene expression of negative regulators of the LepR pathway in the hypothalamus of mice from the study described in Fig. 2. Both TZP and TZP+FcLep groups significantly downregulated Socs3 compared with the FcLep group (Fig. 5F). Furthermore, the TZP+FcLep group downregulated Shp2, while TZP group downregulated Ptp1b (Fig. S5A-B). In contrast, the expression level of Tcptp remained unchanged (Fig. S5C). Moreover, we investigated the hypothalamic transcript levels of neuropeptides involved in energy regulation. The FcLep group exhibited the highest mRNA expression of Pomc. Both TZP and TZP+FcLep groups showed decreased Pomc mRNA levels compared to the FcLep group, which may reflect a compensatory response for long-term weight loss (Fig. 5G). Correspondingly, we observed upregulated orexigenic Npy mRNA in TZP group. Notably, the TZP+FcLep group significantly downregulated Npy mRNA compared to TZP group (Fig. 5H). Considering the roles of area postrema (AP) and nucleus of the solitary tract (NTS) in regulating appetite and metabolism, we evaluated the mRNA level of four key receptors in the hindbrain: Glp1r, Gfral, Calcr, and Slc6a2 (*21*). However, no significant change was observed (Fig. S5D-G).

### Tirzepatide and leptin engaged different cellular targets to exert synergistic effects

A validation RNA in situ study confirmed that LepR and GLP-1R are co-expressed in both POMC and AgRP neurons, with colocalization observed in approximately 20% POMC neurons and 5% of AgRP neurons, respectively (Fig. 6A–D). We next asked whether GLP-1R activation can potentiate LepR signaling in a heterologous system in vitro. To address this question, we transiently expressed GLP-1R and LepR individually, or both receptors simultaneously in HEK293 cells and measured pSTAT3 response to leptin in the presence and absence of GLP-1R agonist Exendin-4 (EX4). Leptin treatment dose-dependently increased pSTAT3 signal in HEK293 cells expressing LepR (Fig. 6E) or LepR + GLP-1R (Fig. 6F), but not in HEK293 cells expressing GLP-1R alone (Fig. 6G). Importantly, EX4 treatment (1nM or 10nM) had no impact on pSTAT3 response to leptin in cells expressing either LepR alone or LepR + GLP-1R (Fig. 6E-F). As a positive control, we confirmed that 10nM EX4 treatment stimulated strong cAMP response in HEK293 cells co-expressing LepR and GLP-1R (Fig. S6). Together, these results suggest that GLP-1 and leptin independently act on their cognate receptors and activate distinct downstream intracellular signaling components.

**Fig. 6.**
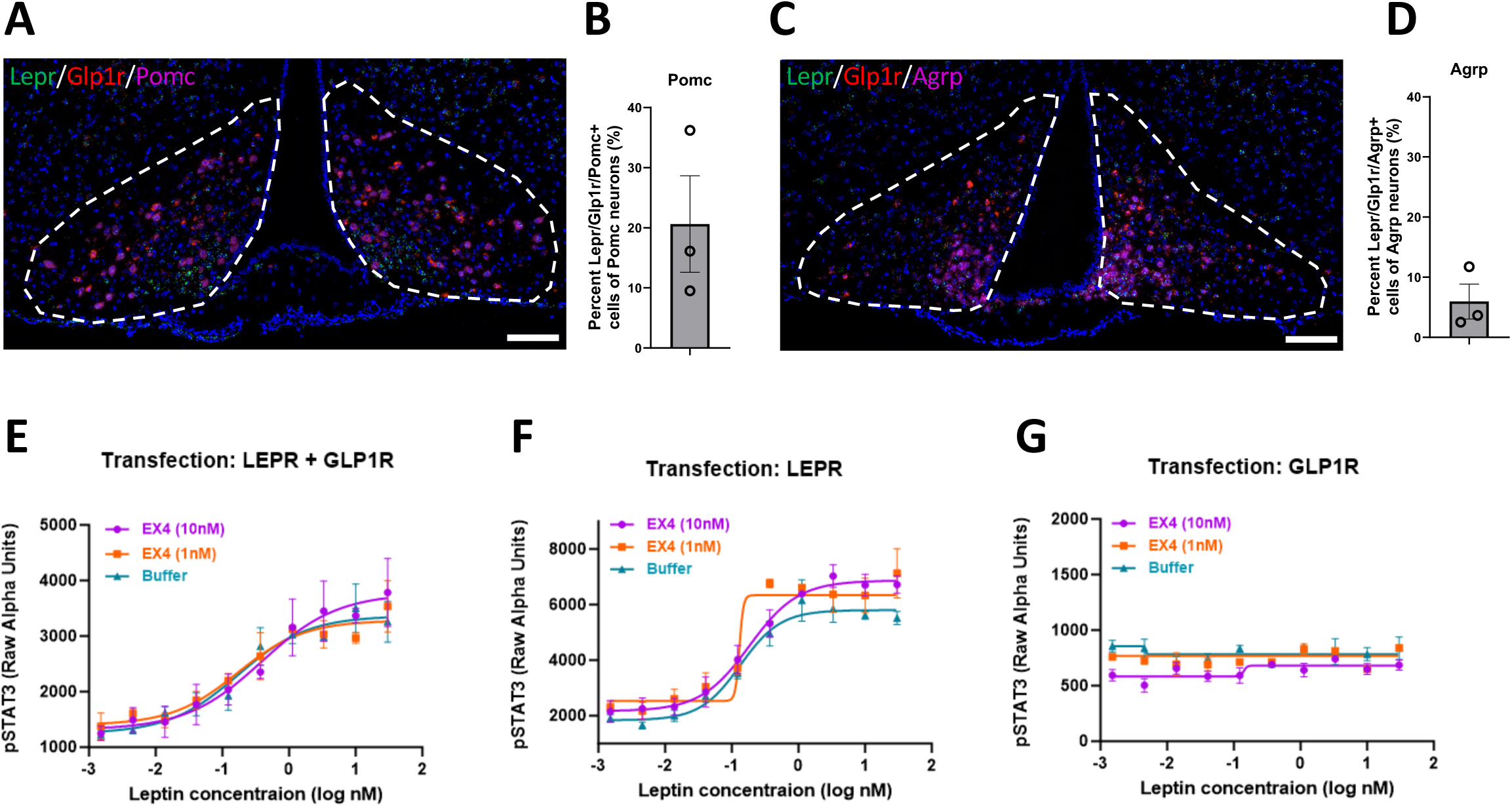
Tirzepatide and leptin engage different intra-cellular pathways. LepR = leptin receptor, GLP-1R = glucagon-like peptide-1 receptor, pSTAT3 = phosphorylated STAT3, Pomc = pro-opiomelanocortin, Agrp = Agouti-related peptide, EX4 = Exendin4. (A) Representative in situ images of Lepr, Glp1r, and Pomc positive cells in arcuate nucleus of hypothalamus. Scale bar: 100um. (B) Bar graph quantification of the overlapping Lepr/Glp1r/Pomc out of total Pomc (%). n = 3. (C) Representative in situ images of Lepr, Glp1r, and Agrp positive cells in arcuate nucleus of hypothalamus. Scale bar: 100um. (D) Bar graph quantification of the overlapping Lepr/Glp1r/Agrp out of total Agrp (%). n = 3. (E-G) The dose-response curves of pSTAT3 induction by leptin co-treated with fixed concentration of EX4 (0 nM, 1 nM, 10 nM, respectively) in HEK293 cells transfected with human LepR/GLP1R (E), LepR alone (F), or GLP1R alone (G). N = 3 for each leptin dose level. Data are displayed as means ± SEM.

### Tirzepatide synergized with leptin to promote hypothalamic POMC neuronal firing

To further investigate the action and mechanism of tirzepatide and leptin combination on POMC neurons, we examined the electrophysiological properties of POMC cells in ARH using acute brain slices from POMC-EGFP transgenic mice (Fig. 7A). Under basal conditions, 6 of 15 recorded cells exhibited burst firing, 5 cells showed spontaneous firing without bursts, and 5 cells were silent at rest, indicating the heterogeneous nature of POMC neuron activity (data not shown). The recorded neurons were then divided into two groups: one was initially treated with recombinant Leptin (Lep, 20 nM), and the other with TZP (10 nM), followed by the respective second treatment (TZP or Lep) (Fig. 7B). TZP and Lep synergistically increased the frequency of action potentials, with no significant changes observed in resting membrane potential or action potential amplitude, indicating increased POMC neuronal firing (Fig. 7C-E).

**Fig. 7.**
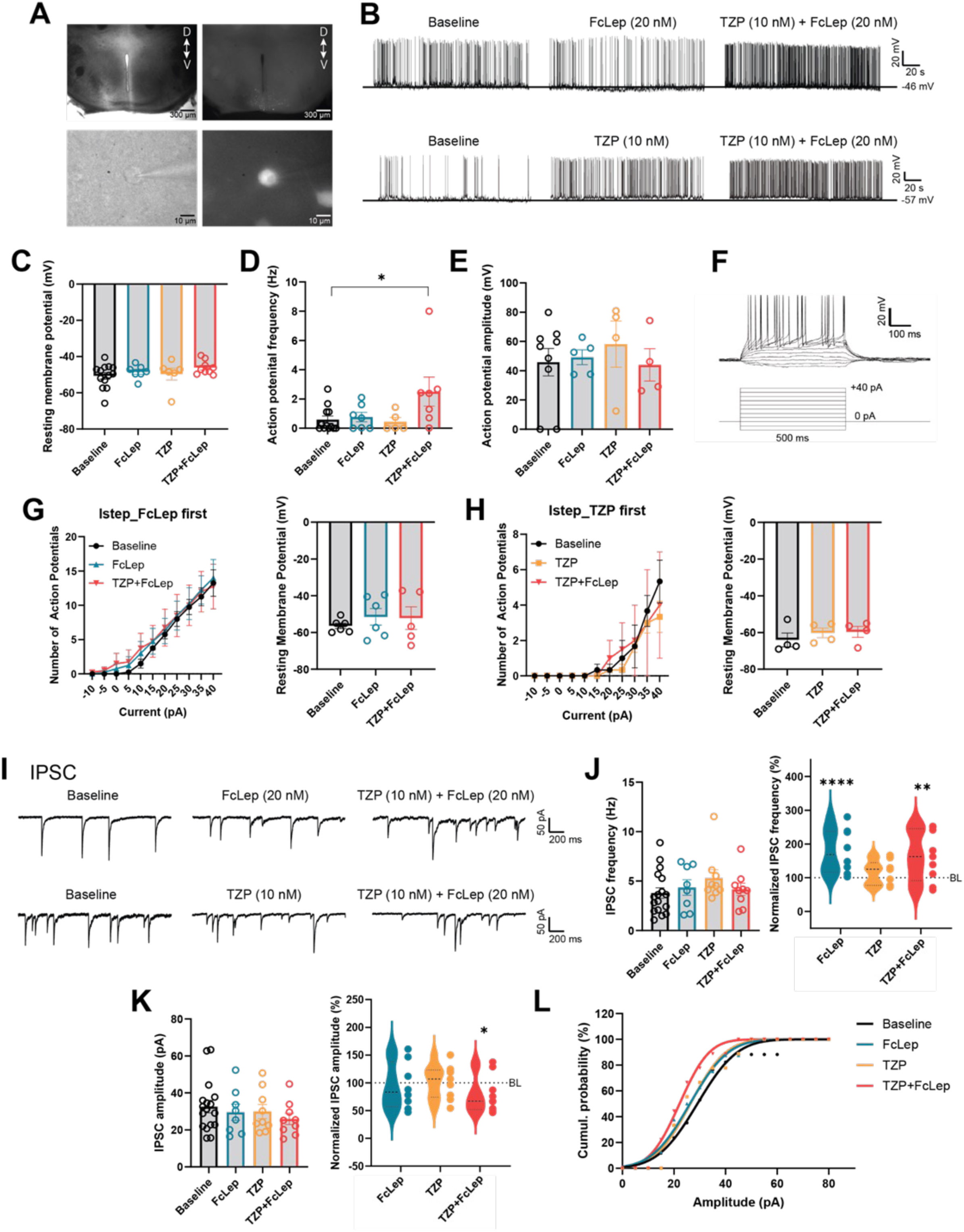
Tirzepatide synergized with leptin to promote hypothalamic POMC neuronal firing. FcLep = mouse IgG1-Fc and human leptin fusion recombinant protein, TZP = tirzepatide, IPSC = inhibitory postsynaptic currents, BL = baseline. (A) Whole-cell patch clamp recording in POMC-GFP neurons, which were identified by an LED illumination system. (B) Representative traces for two groups: one was initially treated with FcLep (20 nM), and the other with TZP (10 nM), followed by the respective second treatment (TZP or FcLep). (C-E) Spontaneous POMC neuron resting membrane potential (mV), firing frequency (Hz), and amplitude (mV). n = 4-13 per group. One-way ANOVA. *p < 0.05. (F) Representative traces for intrinsic excitatory injecting current steps recordings. (G) The effect of FcLep and Lep+FcTZP treatment on the number of action potential and resting membrane potential (mV) in POMC neurons under current injection. n = 5-6 per group. (H) The effect of TZP and TZP+FcLep treatment on the number of action potential and resting membrane potential (mV) in POMC neurons under current injection. n = 4 per group. (I) Representative IPSC traces of POMC neurons under tirzepatide and leptin combo treatment with FcLep first or TZP first. (J) Raw and quantified IPSC frequency of POMC neurons under TZP+FcLep treatment. n = 8-17 per group. Unpaired two-tailed Student’s t test. ^∗∗^p < 0.01, ^∗∗∗∗^p < 0.0001, compared to baseline. (K) Raw and quantified IPSC amplitude of POMC neurons under TZP+FcLep treatment. n = 8-17 per group. Unpaired two-tailed Student’s t test. *p < 0.05, compared to baseline. (L) Cumulative probability of IPSC amplitude. n = 8-17 per group. Data are displayed as means ± SEM.

To further understand the mechanism underlying the increased action potential frequency, we first examined the response of POMC neurons to depolarizing current steps (Fig. 7F). Analysis of action potentials evoked by a 40-pA current injection revealed that POMC neurons fired similarly under both TZP+Lep combination treatment and single reagent (Lep or TZP) treatment, suggesting that TZP+Lep does not alter the activity of voltage-gated ion channels (Fig. 7G-H). Next, we assessed the effect of TZP+Lep on inhibitory postsynaptic currents (IPSCs) mediated by GABAergic neurotransmitters (Fig. 7I). The combination treatment increased IPSC frequency normalized by baseline, potentially driven by leptin (Fig. 7J). In contrast, both normalized IPSC amplitude and size distribution were significantly reduced under TZP+Lep treatment (Fig. 7K-L). These findings suggest a change in postsynaptic input. Together, electrophysiological data in anorexigenic POMC neurons is consistent with the physiological effects observed in DIO mice, such as increased satiety and enhanced energy expenditure, following tirzepatide and leptin combination treatment.

## DISCUSSION

Our post hoc analysis of a clinical study showed higher baseline leptin levels were associated with greater weight loss in patients treated with various doses of tirzepatide, leading to the hypothesis that leptin and tirzepatide could synergize to achieve stronger effects on weight loss and better maintenance of metabolic homeostasis. We treated the DIO mice with tirzepatide, leptin or their combination. Our results showed tirzepatide and leptin synergistically promoted weight loss through reduced food intake and increased energy expenditure, which led to improved insulin sensitivity and metabolism in hepatic and adipose tissues. Tirzepatide sensitizes leptin signaling in hypothalamic GLP-1R expressing and POMC neurons. Tirzepatide and leptin synergistically increase POMC neuronal firing by decreasing the postsynaptic inhibitory input onto these neurons.

Tirzepatide and leptin combination treatment yielded greater effects on weight loss and feeding suppression than each treatment alone. At the end of the 21-day treatment time course, mice treated with tirzepatide and leptin combination had the lowest liver weight and highest hepatic Irs2 expression compared to the control and single treatment groups. Irs2 is an insulin signaling adaptor and essential for suppressing hepatic glucose production (HGP) and promoting glucose uptake in the liver (*22*). The TZP+FcLep treatment group and the TZP group had the lowest hepatic gluconeogenic and lipogenic gene expression among all groups. Our results indicate tirzepatide and leptin combination treatment improved hepatic insulin sensitivity.

Mice treated with tirzepatide and leptin combination had the lowest white and brown adipose tissue weight at the end of the 21-day treatment time course. The expression of the thermogenic genes, including Ucp1 and Elovl3, was the highest in the brown adipose tissue in the TZP+FcLep group compared to the control and single peptide treatment groups. GLP-1R agonism promotes thermogenesis and increases Ucp1 expression in the brown adipose tissue (*23*), though the effect of leptin on thermogenesis is still debatable (*24–27*). Our results indicate tirzepatide and leptin synergistically increase thermogenesis in the brown adipose tissue, which could contribute to the weight loss effects. These mice were group housed in cages at room temperature, which is below their thermoneutral zone and potentially affects the metabolic outcome. However, our second 21-day time course study addressed this concern. We repeated the experiment using mice individually housed in metabolic cages under thermoneutrality and replicated the findings of weight loss and reduced feeding. Moreover, we showed that mice treated with tirzepatide and leptin combination had the highest oxygen consumption and energy expenditure among the four groups during week 2 despite of their low body weight. Together, these data strongly support our hypothesis that tirzepatide and leptin synergize to promote weight loss through reduced feeding and increased energy expenditure.

We tested the effects of tirzepatide on leptin sensitization in DIO mice by examining the acute responses to leptin in the brains of mice treated with vehicle or tirzepatide. Our quantified results showed 6-day tirzepatide treatment significantly increased the hypothalamic POMC and GLP-1R neuronal response to a bolus leptin injection. As reduced feeding is often associated with increased leptin sensitivity, we include the pair-feeding (PF) group as a control. Pair-feeding also increased leptin sensitivity in POMC neurons. However, only tirzepatide treatment increased leptin sensitivity in hypothalamic GLP-1R neurons, indicating this is a drug-specific effect, not just a secondary effect to reduced feeding.

Studies from other groups tested the various approaches of combining long-term adiposity signal (leptin) and short-term satiation signal (incretin hormones) for weight loss. For instance, both preclinical and clinical evidence suggest that the combination of amylin and leptin results in greater weight loss than amylin alone (*28, 29*). Addition of leptin to GLP-1/glucagon co-agonism leads to greater weight loss and reductions in food intake in DIO mice compared to GLP-1/glucagon co-agonism monotherapy (*30*). Recent studies identified the GLP-1R/LepR-coexpressing neurons in the dorsomedial hypothalamic nucleus (DMH) (*31*). Polex-Wolf et al. developed a dual GLP1R/LepR agonist and evaluated its physiological efficacy in several mouse models, though the synergistic effect of the GLP-1 and leptin remains unclear due to technical issues (*17*). In our study, we test if leptin and GLP-1 signaling pathways directly interact to increase leptin signaling using in vitro cultured cells expressing both GLP-1R and LepR. Addition of GLP-1 resulted in no change of the leptin dose-response curves, suggesting simultaneously activating the lepR and GLP-1R signaling pathways cannot augment leptin signaling.

Previous studies indicated that the loss of leptin sensing in the melanocortin circuits likely contributed to the pathology of leptin resistance (*32*). Human cerebrospinal fluid (CSF) POMC, but not AGRP, peptide showed significant negative correlation with CSF leptin, plasma leptin, and adiposity (*33*). Our data showed less Pomc and Scos3 transcripts in the hypothamali of the mice chronically treated with tirzepatide and leptin combination or tirzepatide alone, consistent with increased leptin sensitivity in these mice. We measured the acute response of POMC neurons to leptin, tirzepatide, or combination treatment. The increased POMC neuronal firing response with tirzepatide and leptin combination is not a result of altered POMC neuron intrinsic firing properties but likely caused by decreased inhibitory post-synaptic input from other neurons. We hypothesize that tirzepatide and leptin act through different neuronal targets and neurocircuitries, part of which converge on POMC neurons. We showed a small fraction of POMC neurons is LepR and GLP-1R double positive. This is consistent with the previous report of LepR and GLP-1R-expressing POMC neurons represent largely nonoverlapping subpopulations (*34*).

Our findings support the notion that the effect of tirzepatide and leptin on POMC neurons is indirect. However, a limitation of our study is we have not elucidated the identity of the source of the inhibitory postsynaptic projects to the POMC neurons. A candidate could be NPY/AgRP neurons, as the GABAergic nerve terminals of these neurons synapse on the POMC neurons (*35*). Nor have we identified the neural circuitries of tirzepatide and leptin engage and converge on to regulate feeding and peripheral metabolism. Here, we focused on the actions of tirzepatide and leptin in hypothalamic neurons. The neural circuitries of GLP-1R and lepR extend well beyond the hypothalamus (*1, 36–40*). To fully address this intriguing question is beyond the scope of the current study and warrants future investigation. Studies from other groups reported both direct and indirect effects of liraglutide on POMC neurons (*41*). The different findings could be attributed to the different type and doses of drugs used. Notably, we used much lower doses of tirzepatide (10nM vs 1uM liraglutide(*41*)) and leptin (20nM vs 100nM(*41*)). Tirzepatide is a GLP-1R and GIPR co-agonist. Both GIPR agonism and antagonism were used for weight loss strategies through different mechanisms of action (*42*). GIP was reported to impair neural leptin sensitivity in DIO mice (*43*), while loss of GIPR in LepR cells did not affect GIP:GLP-1 co-agonism effects on weight control (*44*). We have yet distinguished the molecular actions of tirzepatide through GLP-1R versus GIPR, which remains a limitation of the current study.

Biomarker analysis using human blood samples from tirzepatide clinical trials revealed that higher circulating leptin baseline levels correlate with greater weight loss. This was consistent with both female and male patients. Our studies showed the combination of tirzepatide and leptin caused synergistic effects on weight loss and feeding suppression in DIO mice housed at ambient temperature or under thermoneutrality. Weight loss in patients with obesity is often associated with increased hunger, reduced energy expenditure, and other adaptive changes in central and autonomic nervous system. Weight loss reduces fat mass and decreases circulating leptin. Leptin administration in patients reverses weight loss-induced changes in energy expenditure, autonomic, neuroendocrine adaptations, and regional neural activity stimulated with visual food cue (*45–47*). Our results showed the combination treatment group had better feeding suppression and higher energy expenditure during the weight loss time course, suggesting the tirzepatide and leptin combination could be an effective strategy to combat obesity recidivism. A Phase 2 randomized double-blind clinical trial (NCT06373146) is underway to determine if combining tirzepatide with mibavademab (a LepR agonist) will result in more weight loss than tirzepatide alone.

In conclusion, our results highlight the potential of leptin and incretin hormones combination therapies, particularly tirzepatide, in promoting weight loss and maintenance. Future research to address the molecular and neuronal bases of similar combination therapies will advance our knowledge in neuroendocrine regulation of feeding behavior and overall metabolism.

## Acknowledgments

Drs. Ai and Ren are the guarantors of this work. We thank the technical support from scientists at GUBRA, the undergraduate students at Indiana University, and the islets core at IU Center for Diabetes and Metabolic Diseases.

## Author contributions

X.S., M.A., and H. R. designed, conducted experiments, analyzed data, wrote and revised the manuscript. J.M.W. and Y.L. conducted and analyzed the post hoc human clinical data. B.A.D. and R.J.S. designed and performed indirect calorimetry experiments. Y.Y. and P.L.S. designed and performed electrophysiology experiments. M.S. and M.A. performed the molecular pharmacology experiments. B.Z., H.C.R and P.F.C provided research reagents and resources. H.R. and M.A. conceived and supervised the study. All authors reviewed and approved the manuscript.

---

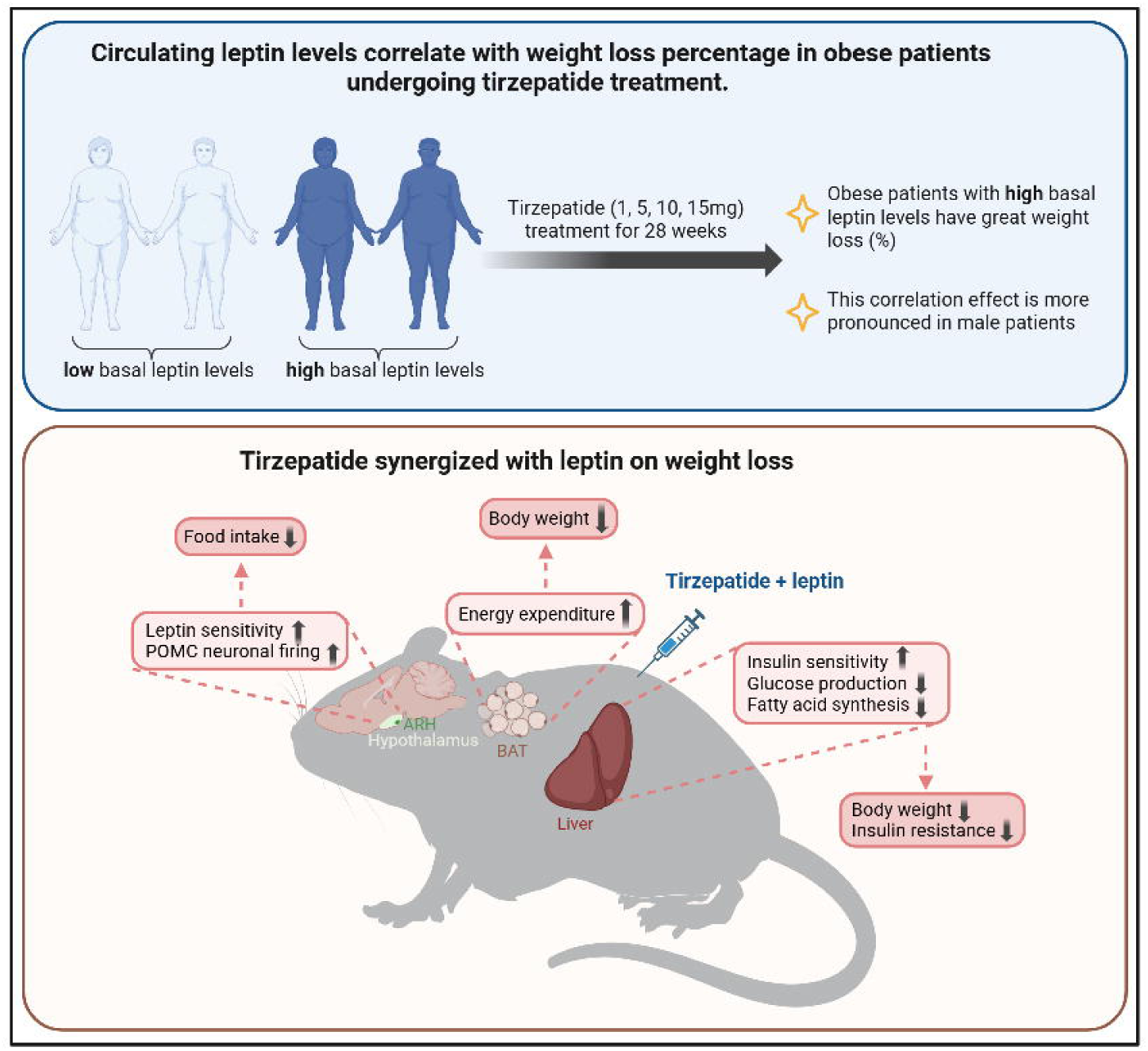

