## Supplementary Figures and Tables for "Tirzepatide Synergizes with Leptin on Weight Loss and Restoring Metabolic Homeostasis in Diet-induced Obesity Model": Supplementary Materials.pdf

[4]Department of Pharmacology & Toxicology, Indiana University School of Medicine; Indianapolis, IN, USA.

Postal address: 307 East Merrill St., Indianapolis, IN 46225

### MATERIALS AND METHODS

**Protein sequence of Fc-Leptin:**

VPRDSGCKPCICTVPEVSSVFIFPPKPKDVLTITLTPKVTCVVVDISKDDPEVQFSWFVDDVEVHT
AQTQPREEQFNSTFRSVSELPIMHQDWLNGKEFKCRVNSAAFPAPIEKTISKTKGRPKAPQVYTIP
PPKEQMAKDKVSLTCMITDFFPEDITVEWQWNGQPAENYKNTQPIMDTDGSYFVYSKLNVQKS
NWEAGNTFTCSVLHEGLHNHHTEKSLSHSPGGGGSVPIQKVQDDTKTLIKTIVTRINDISHTQSVS
SKQKVTGLDFIPGLHPILTLSKMDQTLAVYQQILTSMPSRNVIQISNDLENLRDLLHVLAFSKSCH
LPWASGLETLDSLGGVLEASGYSTEVVALSRLQGSLQDMLWQLDLSPGC

Red fonts: mouse IgG1-Fc

Blue fonts: linker amino acid sequence

Back fonts: human leptin sequence without signal peptide

**Fig. S1**

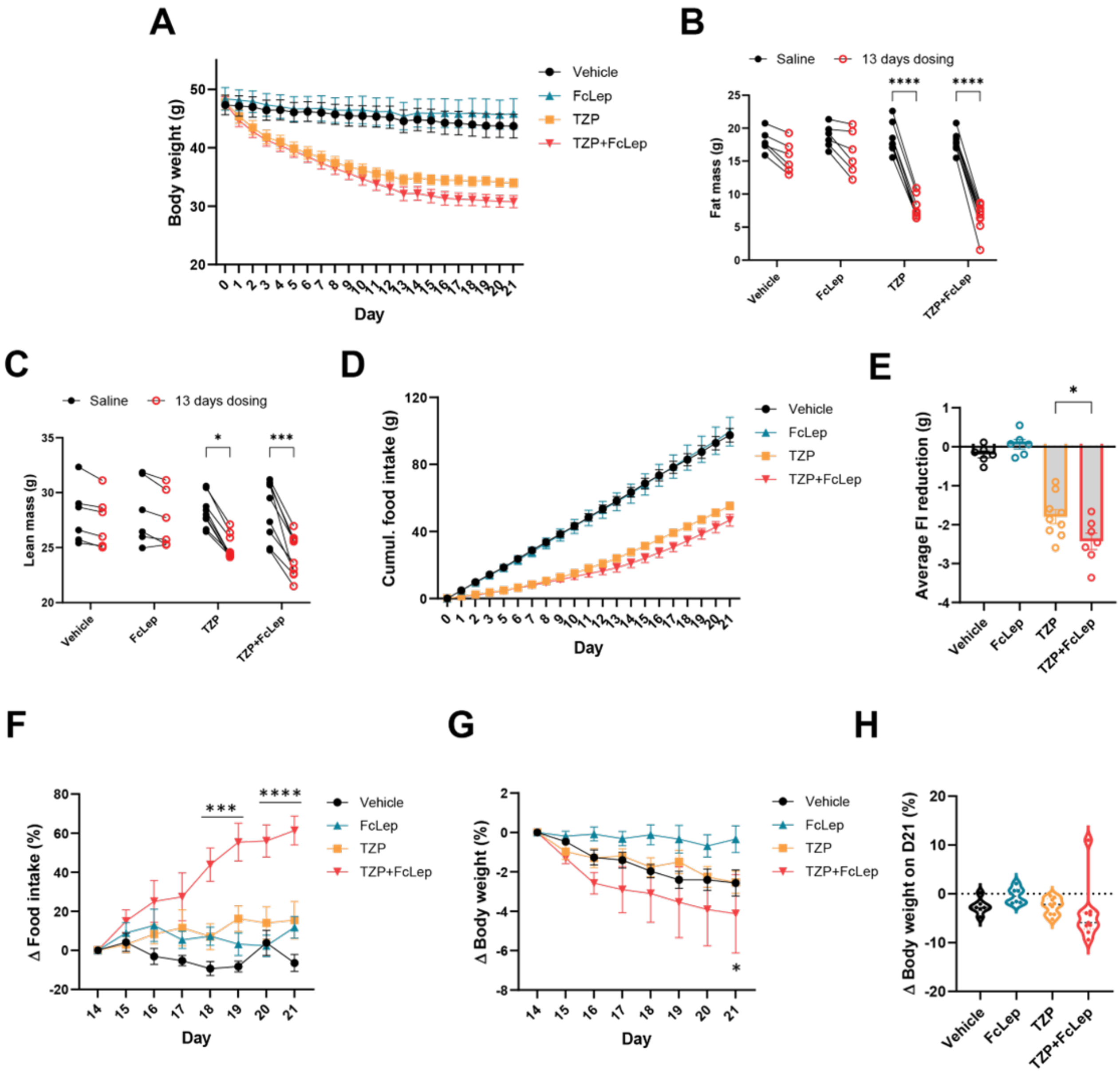


**Tirzepatide synergized with leptin on weight loss**

FcLep = IgG1-Fc-leptin fusion protein, TZP = tirzepatide, FI = food intake. (A) Body weights (g) throughout 21 days of dosing treatment. (B-C) Fat mass and lean mass (g) one day before and after 13 days of dosing treatment. Two-way ANOVA. (D) Cumulative food intake throughout 21 days of dosing. (E) Average 24 h food intake reduction (g). One-way ANOVA. (F-G) Normalized food intake reduction (%) and body weight reduction (%) during the last week (day 14-21). Two-way ANOVA. (H) Normalized body weight reduction (%) on day 21. n = 6-9. $^{*}p < 0.05$, $^{***}p < 0.001$, $^{****}p < 0.0001$. Data are displayed as means ± SEM.

**Fig. S2**

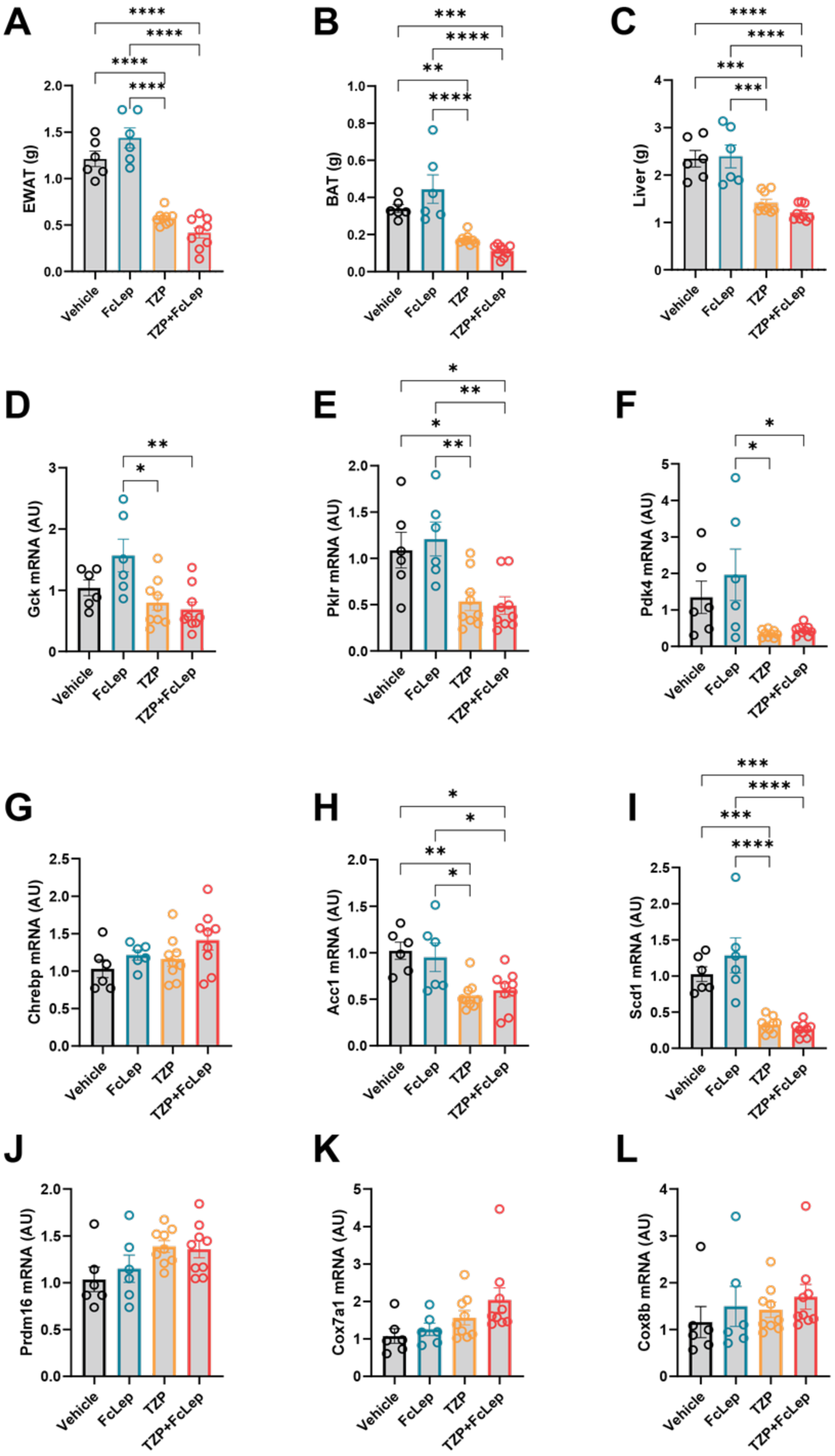
A
EWAT (g)
B
BAT (g)
C
Liver (g)
D
Gck mRNA (AU)
E
Pklr mRNA (AU)
F
Pdk4 mRNA (AU)
G
Chrebp mRNA (AU)
H
Acc1 mRNA (AU)
I
Scd1 mRNA (AU)
J
Prdm16 mRNA (AU)
K
Cox7a1 mRNA (AU)
L
Cox8b mRNA (AU)
Vehicle
FcLep
TZP
TZP+FcLep

**Tissue weights and altered expression of metabolic genes**

FcLep = IgG1-Fc-leptin fusion protein, TZP = tirzepatide, EWAT = epididymal white adipose tissue, BAT = brown adipose tissue. (A-C) Tissue weights (g) of EWAT, BAT, and liver. (D-I) qPCR analysis of Gck, Pklr, Pdk4, and Chrebp, Acc1, and Scd1 in liver tissue. (J-L) qPCR analysis of Prdm16, Cox7a1, and Cox8b in BAT tissue. One-way ANOVA was performed. n = 6-9. *$p < 0.05$, **$p < 0.01$, ***$p < 0.001$, ****$p < 0.0001$. Data are displayed as means ± SEM.

**Fig. S3**

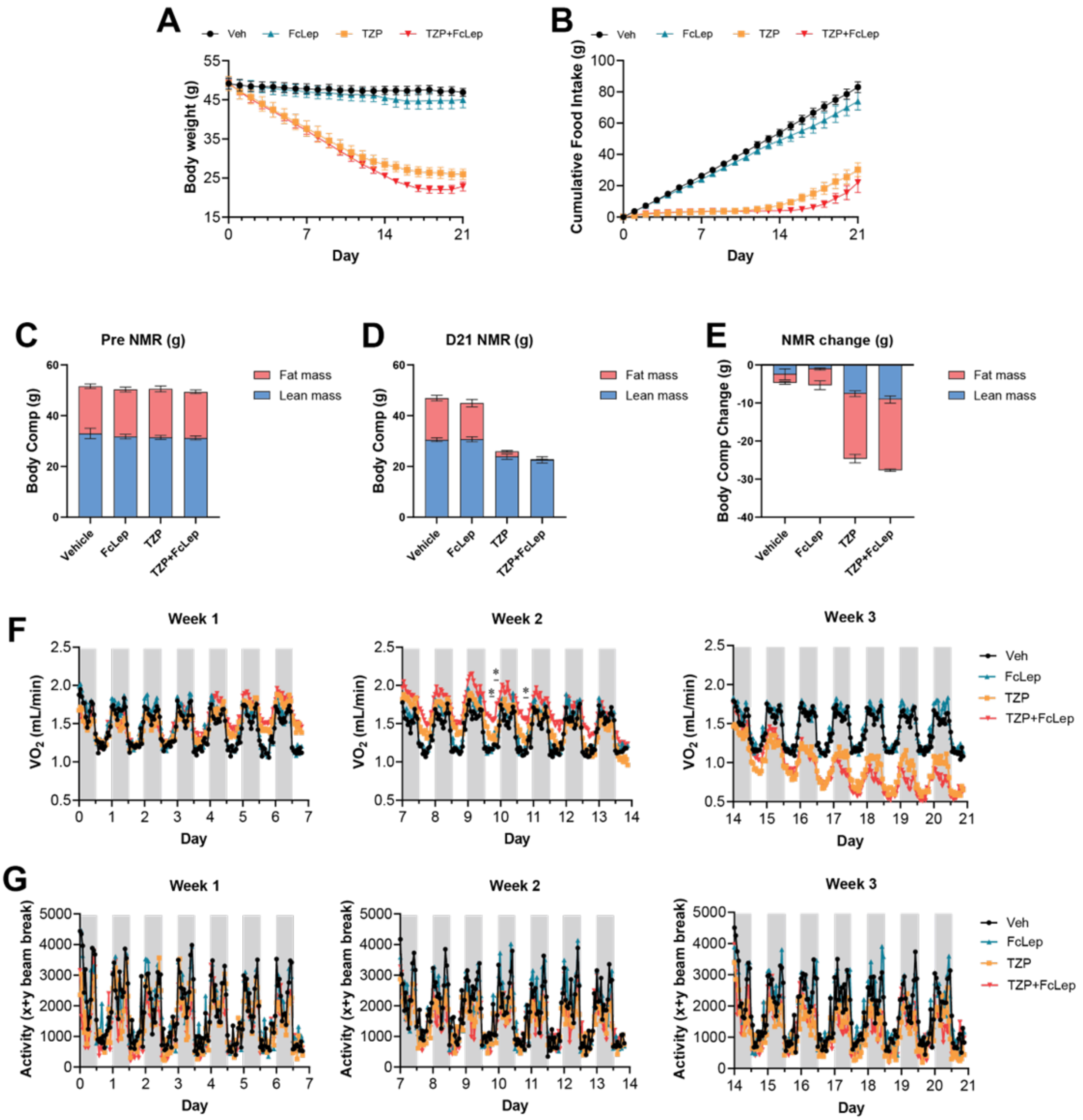


**Increased energy expenditure in DIO mice treated with TZP+FcLep compared to TZP alone**

FcLep = IgG1-Fc-leptin fusion protein, TZP = tirzepatide, NMR = nuclear magnetic resonance, EE = energy expenditure. (A) Body weights (g) throughout 21 days of dosing (vehicle, FcLep, TZP, TZP+FcLep) at thermoneutrality. (B) Cumulative food intake (g) throughout 21 days of dosing at thermoneutrality. (C-D) Fat mass and lean mass (g) of DIO mice before 3-day and after 21-day of dosing treatment. (E) The change of fat mass and lean mass (g) after 3-week treatment. (F) Rhythms oxygen consumption during 3-week treatment. Unpaired two-tailed Student's t test was performed. $^{*}p < 0.05$, statistically significant at three consecutive time points, compared to Tzp. (G) The activity (x+y beam break) of DIO mice during 3-week treatment. n = 6 per group. Data are displayed as means ± SEM.

**Fig. S4**

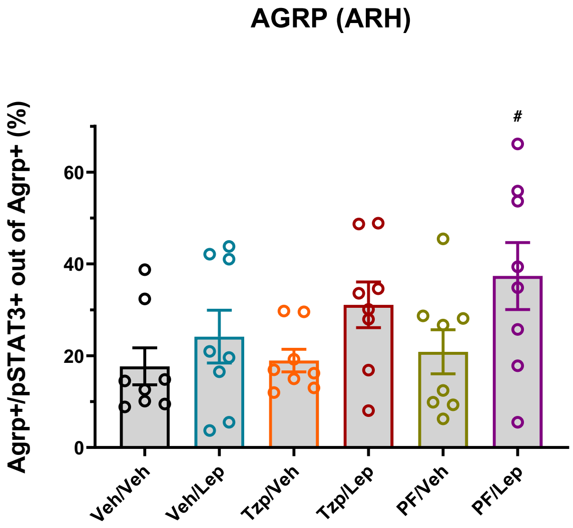


**Quantification of pSTAT3 response in AgRP+ neurons**

Veh = vehicle, Lep = leptin (LY355101), TZP = tirzepatide, PF = pair-feeding, ARH = arcuate nucleus of hypothalamus, AgRP = agouti-related protein, pSTAT3 = phosphorylated STAT3. n = 8. Dunnett's test one-factor linear model. No statistical significance among the treatment groups.

**Fig. S5**

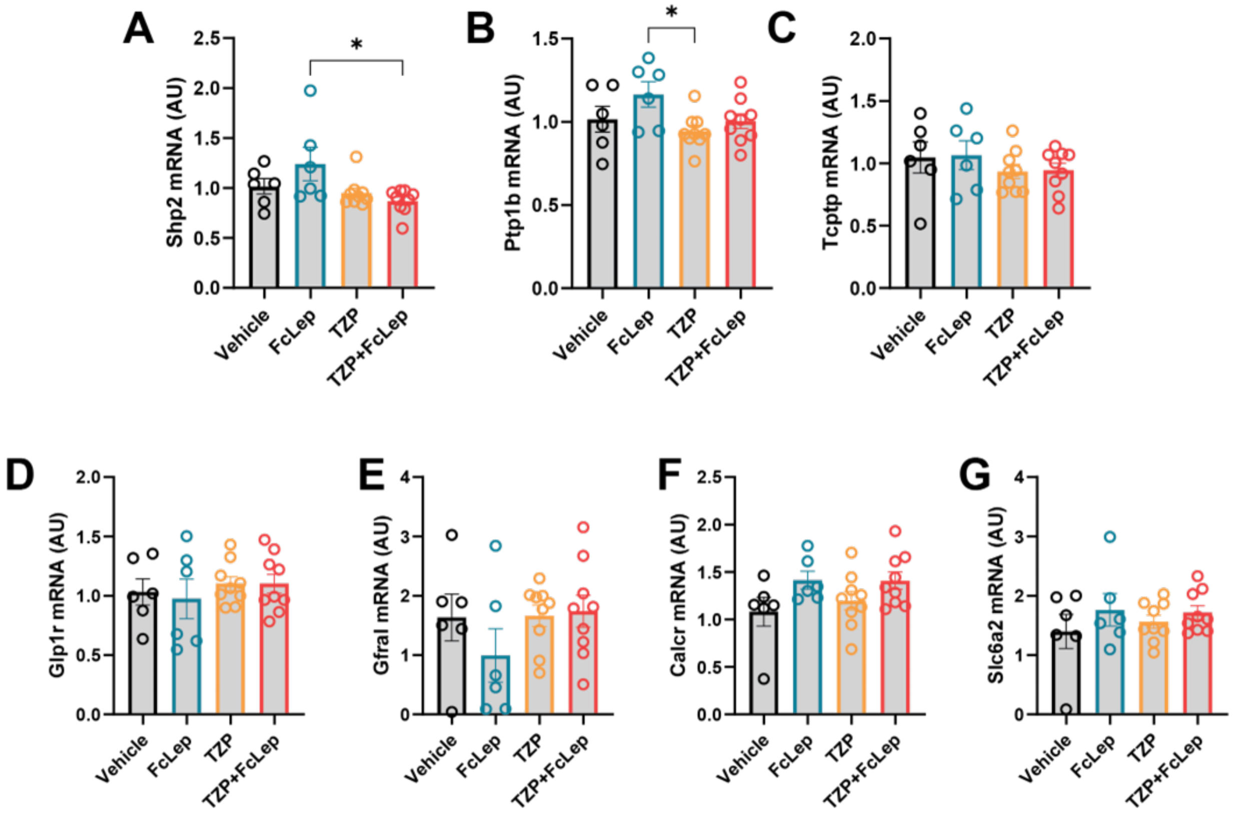


**Gene expression levels in hypothalamus and hindbrain**

FcLep = IgG1-Fc-leptin fusion protein, TZP = tirzepatide. (A-C) qPCR analysis of Shp2, Ptp1b, and Tcptp in hypothalamus. n = 6-9. *$p < 0.05$. One-way ANOVA. (D-G) qPCR analysis of Glp1r, Gfral, Calcr, and Slc6a2 in hindbrain. Data are displayed as means ± SEM.

**Fig. S6**

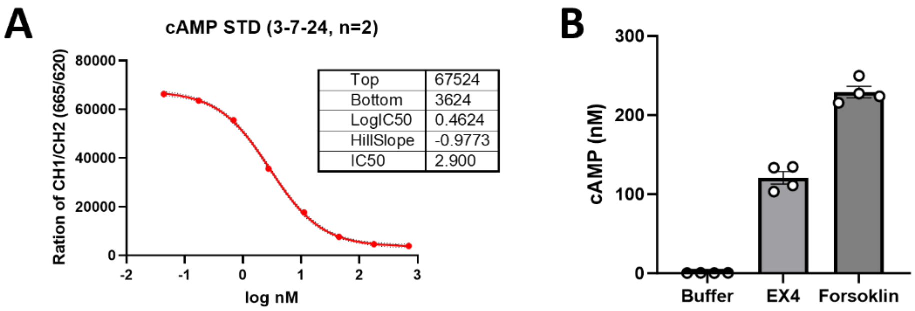


**Exendin-4 induced cAMP response in GLP1R-expressing HEK293 cells**

cAMP = cyclic adenosine monophosphate, EX4 = Exendin-4. (A) Standard curve of cAMP assays in HEK293 cells. (B) The cAMP induced by EX4 (10nM) and Forskolin (1uM) in HEK293 cells expressing GLP1R. (n = 4 in each group). Data are displayed as means ± SEM.

**Table S1**

| Genes | Forward primer sequence (5'->3') | Reverse primer sequence (5'->3') |
|---|---|---|
| Irs2 | AGTGATGGGACAGGAAGTCG | TCCAGAACGGCCTCAACTAT |
| G6pc | GCCAACCGGGCTGGACTCAC | AGGCACGGAGCTGTTGCTGT |
| Srebp1c | GAAGCTGTCGGGGTAGCGTCT | CTCTCAGGAGAGTTGGCACCTG |
| Fas | TGAGCTGGGTTAGGGTAGGA | CTGACTCGGCTACTGACACG |
| Gck | GGCCTCCGGAGCAGAAGGGA | ACACATGCGCCCCTCATCGC |
| Pklr | CTTGCTCTACCGTGAGCCTC | ACCACAATCACCAGATCACC |
| Pdk4 | AGCTGCTGGACTTTGGTTCAGAAAAT | TGGGCTCTTCTCATGGAACTCCACC |
| Chrebp | GGCTGTGCAAGCTGGCGGTA | CACCGGGGTGCCCATCACAC |
| Acc1 | GGACAGACTGATCGCAGAGAAAG | GCTGTTCCTCAGGCTCACAT |
| Scd1 | CTCCTGCTGATGTGCTTCAT | AGGGTGCTAACGAACAGGCT |
| Ucp1 | ACTGCCACACCTCCAGTCATT | CTTTGCCTCACTCAGGATTGG |
| Pgc1a | CCCTGCCATTGTTAAGACC | TGCTGCTGTTCCTGTTTTC |
| Elovl3 | ATGCAACCCTATGACTTCGAG | ACGATGAGCAACAGATAGACG |
| Cidea | TGCTCTTCTGTATCGCCCAGT | GCCGTGTTAAGGAATCTGCTG |
| Prdm16 | CAGCACGGTGAAGCCATTC | GCGTGCATCCGCTTGTG |
| Cox7a1 | GCTCTGGTCCGGTCTTTTAG | CTTTCAAGTGTACTGGGAGGTC |
| Cox8b | GAACCATGAAGCCAACGACT | GCGAAGTTCACAGTGGTTCC |
| Pomc | TAGATGTGTGGAGCTGGTGC | TTTTCAGTCAGGGGCTGTTC |
| Npy | TAGGTAACAAGCGAATGGGG | TGTCTCAGGGCTGGATCTCT |
| Ptp1b | GGAACAGGTACCGAGATGTCA | AGTCATTATCTTCCTGGGTGCAATT |
| Tcptp | AGGGCTTCCTTCTAAGG | GTTTCATCTGCTGCACCTTCTGAG |
| Shp2 | AGCTGGCTGAGACCACAGAT | CCTGTTGCTGGAGCGTCT |
| Socs3 | CACCTGGACTCCTATGAGAAAGTG | GAGCATCATACTGATCCAGGAACT |
| Gfral | CCCCACTTGCCTCAGTGTAA | ATGCACACGTGTTCTTCAGC |
| Calcr | TGCATTCCCGGGATACACAG | AGGAACGCAGACTTCACTGG |
| Slc6a2 | AGTTTGTGCAAACGGGCG | GCAATGATGAGGAACAGCGTG |
| Glp1r | GTCATCGCTTCAGCCATCCT | GCAGCCGTGCTATACATCCA |
